# Broken symmetry in the human BK channel

**DOI:** 10.1101/494385

**Authors:** Lige Tonggu, Liguo Wang

## Abstract

Voltage-gated and ligand-modulated ion channels play critical roles in excitable cells. To understand the interplay among voltage-sensing, ligand-binding and channel opening, the structures of ion channels in various functional states need to be determined. Here, the “random spherically constrained” (RSC) single-particle cryo-EM method was employed to study the human large conductance voltage- and calcium-activated potassium (hBK or hSlo1) channels reconstituted into liposomes. The hBK structure has been determined at 3.5 Å resolution in the absence of Ca^2+^. Instead of the common four-fold symmetry observed in ligand-modulated ion channels, a two-fold symmetry was observed in hBK. Two opposing subunits in the Ca^2+^ sensing gating ring rotate around the center of each subunit, which results in the movement of the assembly and flexible interfaces and Ca^2+^ binding sites. Despite the significant movement, the local conformation of the assembly interfaces and Ca^2+^ binding sites remains the same among the four subunits.

## INTRODUCTION

The large conductance voltage- and calcium-activated potassium (BK) channel belongs to the six-transmembrane-segment (6TM) ion channel family. It can be found in many cells and functions as a feedback regulator of membrane potential and thereby Ca^2+^ influx (Cui et al., 2009; Gribkoff et al., 2001; Horrigan and Aldrich, 2002). Like other members of the 6TM ion-channel family, it has voltage-sensor domains (VSDs) which sense the transmembrane potential. The BK channel is also a member of the Slo family, whose α-subunits contain regulator-of-conductance-for-K^+^ (RCK) domains in the large intracellular C-terminal region. The crystal structures of the RCK domain of the *E. coli* K^+^ channel (Jiang et al., 2001) and the gating ring structure formed by eight identical RCK domains in the *Methanobacterium thermoautotrophicum* K^+^ channel MthK (Jiang et al., 2002a, b; Ye et al., 2006) greatly aid the understanding of Ca^2+^-dependent regulation of Slo1 channels. First, these prokaryotic channel structures inspired a series of mutational studies demonstrating that the RCK1 and RCK2 domains in mouse Slo1 (mSlo1) account for Ca^2+^-dependent activation of mSlo1 (Schreiber and Salkoff, 1997; Shi et al., 2002; Sweet and Cox, 2008; Xia et al., 2002; Zhang et al., 2010). Second, these results provide templates for the generation of homology models of the mSlo1 channel gating ring: the four RCK1 and four RCK2 domains from the four subunits in a functional Slo1 channel form a “gating ring”. The high-affinity Ca^2+^ binding site, termed as Ca^2+^ bowl, is located in RCK2 and the mD367 site is in RCK1. The low-affinity Ca^2+^ binding site, E535, resides in the C-terminal lobe of RCK1. These findings were confirmed by the X-ray crystallographic structures of the isolated gating ring of the human Slo1 channel (hSlo1-GR) in the absence of Ca^2+^ (Wu et al., 2010) and that of the zebrafish Slo1 channel (zSlo1-GR) in the presence of Ca^2+^ (Yuan et al., 2012). Recently, the cryo-EM structures of the *Aplysia californica* Slo1 channel (aSlo1) were obtained with and without Ca^2+^, which further confirmed these findings (Hite et al., 2017; Tao et al., 2017). All the gating rings in the structures made of RCK domains (Hite et al., 2017; Jiang et al., 2001; Jiang et al., 2002a, b; Tao et al., 2017; Wu et al., 2010; Yuan et al., 2012; Yuan et al., 2010) have a four-fold symmetry along the central pore (C4). Although the two structures of aSlo1 were determined in both liganded and metal-free states (Hite et al., 2017; Tao et al., 2017), structures of other states will be useful. Functional studies show that the channel has at least four, and probably many more conformational states (Horrigan and Aldrich, 2002) as the channel can be activated by Ca^2+^, voltage, or combinations of the two.

As shown by both structural and functional studies, the lipid membrane environment plays an essential role for the structural integrity and activity of membrane proteins (Gonen et al., 2005; Hilgemann, 2003; Hille et al., 2015; Lee, 2011; Long et al., 2007; Schmidt et al., 2006). Therefore, it is critical to restore the lipid membrane environment of membrane proteins. One method is to use lipid nanodiscs, where the membrane protein resides in a small patch of lipid bilayer encircled by an amphipathic scaffolding protein (Bayburt et al., 2002). The nanodisc method has been employed to study the anthrax toxin pore at 22-Å resolution (Katayama et al., 2010), TRPV1 ion channel in complex with ligands at 3-4 Å resolution (Gao et al., 2016), and other membrane proteins (Autzen et al., 2018; Bayburt et al., 2002; Chen et al., 2017; Dang et al., 2017; Gao et al., 2016; Jackson et al., 2018; Katayama et al., 2010; McGoldrick et al., 2018; Roh et al., 2018; Srivastava et al., 2018; Taylor et al., 2017). An alternative method, called “random spherically constrained” (RSC) single-particle cryo-EM, where the membrane protein is reconstituted into liposomes (a mimic of an empty cell), has also been developed and employed to study the large conductance voltage- and calcium-activated potassium (BK) channels reconstituted into liposomes at 17-Å resolution (Wang and Sigworth, 2009) and a voltage-gated potassium channel Kv1.2 at 12-Å resolution (Jensen et al., 2016). Although both methods restore the lipid environment of membrane proteins, there is a major difference: the RSC method mimics the cell and provides an asymmetric environment (i.e. inside and outside conditions can be varied independently), whereas there is only one environment surrounding the membrane proteins in the nanodisc method. This can be critically important for understanding membrane proteins which are controlled/modulated by asymmetric environment (e.g. applying transmembrane potential for voltage-gated ion channels or voltage-sensitive proteins, applying pressures to mechanosensitive channels).

Here, we employed the RSC method to study the structures of human BK (hBK) channels in the absence of Ca^2+^. By optimizing cryo-EM sample preparation, increasing the data size, employing the direct electron detector camera and correcting the drift between frames with MotionCor2 (Zheng et al., 2016), the resolution has been improved to 3.5 Å. Compared with the commonly observed four-fold symmetry in all known gating ring structures, a broken symmetry was observed in metal-free hBK channels reconstituted into liposomes. Within the tetramer, two opposing subunits in the Ca^2+^ sensing gating ring rotate around the center of mass of each subunit, which results in the movement of the assembly interfaces, flexible interfaces and Ca^2+^ binding sites.

## RESULTS

### Purification and reconstitution of hBK channels into liposomes

Here, we employed the RSC method to study the structures of hBK reconstituted into liposomes. The hBK (KCNMA1) protein was extracted from HEK293 cells stably expressing Flag-tagged human BK proteins, purified by an anti-Flag affinity column followed by a size-exclusion column (Figure S1A), and reconstituted into POPC liposomes with detergent removed by gel filtration. The 125-kDa hBK protein is detected by SDS-PAGE before and after reconstitution (Figure S1B). The reconstitution was adjusted to yield an average protein content at one to two hBK tetramers per 20-nm liposome. As the reconstitution of hBK proteins into liposomes is expected to restore the lipid environment of membrane proteins, the function of the reconstituted proteins was confirmed using ACMA flux assay, which has been employed to assess functions of reconstituted ion channels (Brohawn et al., 2012; Li et al., 2015; Miller and Long, 2012; Su et al., 2016) (Figure 1C-D).

**Figure 1:**
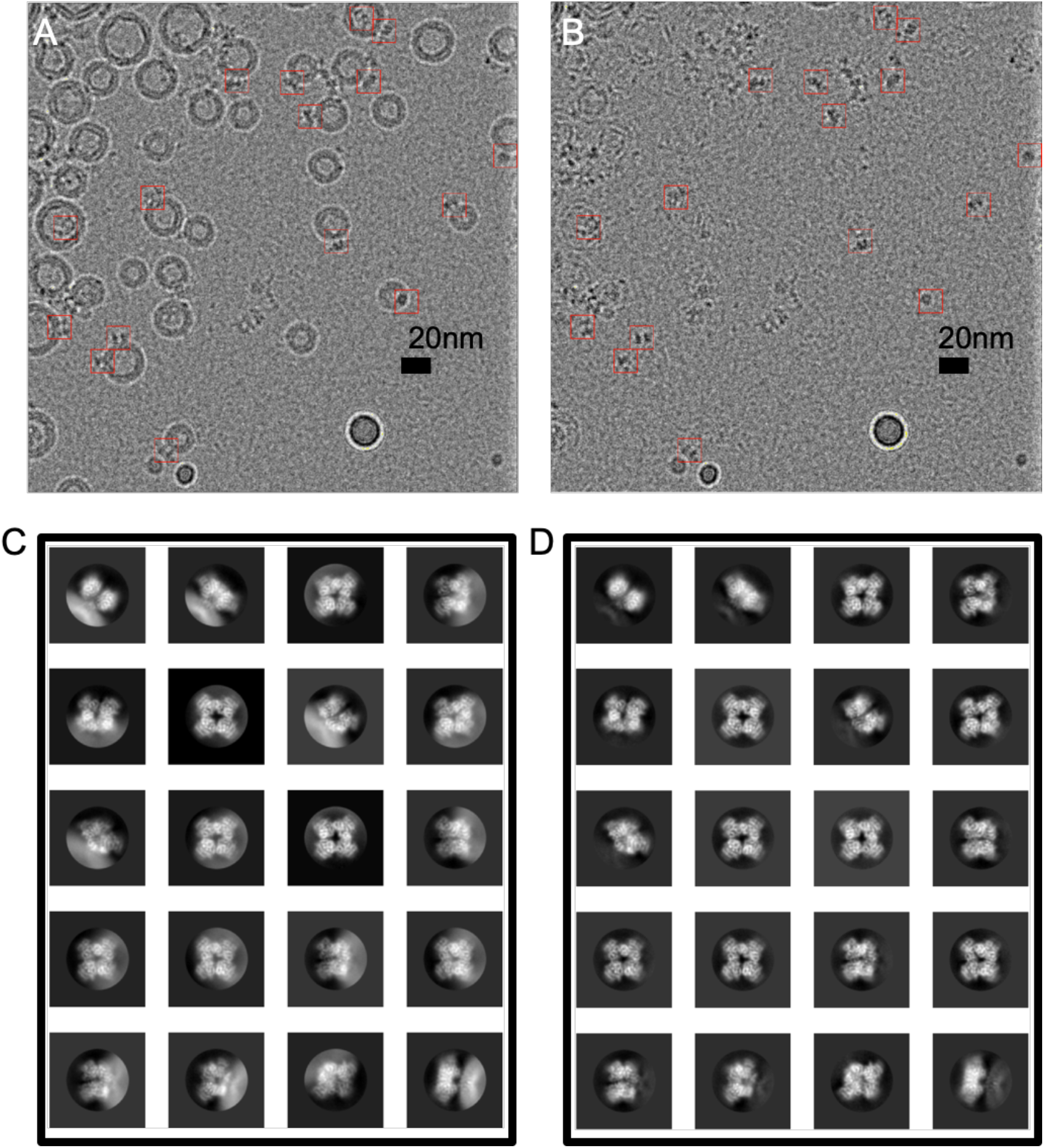
Subtraction of modeled liposome images from an hBK proteoliposome micrograph. (A) A representative cryo-EM image of hBK proteoliposomes at −3.8 μm defocus. (B) Liposomes are subtracted from the image shown A. hBK particles are marked with red boxes (15 nm). (C-D) The largest 20 2D class averages of 122,000 BK particles before (C) and after (D) liposome subtraction. Box size is 27 nm, and the circular mask is 17 nm in diameter. Related to Figure S1.

### Structure of hBK in liposomes

Single-particle reconstruction of unstained cryo-EM specimens typically requires the acquisition of hundreds of thousands of particles. Because the low yield of hBK expressed in HEK293 cells thus the low concentration of proteoliposomes, acquiring such a large number of protein particles in liposomes is challenging. Previously, a 2D streptavidin crystal was used as an affinity surface to tether the biotinylated-lipid-doped proteoliposomes (Wang and Sigworth, 2009). However, empty liposomes bind preferentially to the crystal, reducing further the number of visible protein particles. Therefore, we employed a long-incubation method (a 5-10 min incubation on grids followed by another application of sample before freezing) was used. Twenty-sixty liposomes were observed in each cryo-EM micrograph, which covered 0.16 μm^2^ of specimen (Figure 1A).

Ideally a 3D reconstruction would contain an entire proteoliposome, complete with hBK channel and spherical membrane. Unfortunately, the variability of liposome size precludes the merging of their images; instead we fitted and subtracted a model (Wang et al., 2006) of the membrane contribution to each image (Figure 1B). Protein particles were picked from the resulting liposome-subtracted micrographs and were classified using RELION (Scheres, 2012). Some structural details are visible in the 2D classes (Figure 1C-D). The presence of lipid membrane is confirmed in the 2D classes of protein particles extracted from original cryo-EM micrographs using the orientation information determined with liposome-subtracted protein particle images (Figure 1C-D).

The reconstruction of BK structure was carried out without symmetry. Surprisingly, a two-fold symmetry was observed (Figure 2 and Movie S1). In the structure determined with a C2 symmetry, the central opening of the gating ring defined as the distance between C_α_ atoms of V785 in opposing subunits differs by 2.7 Å along the two diagonals (Figure 2B). The shoulder helix J moves by about 5 Å from two opposing subunits, termed as hBK High, to the other two opposing subunits, termed as hBK Low. (Figure 2C). This movement is comparable to the movement of the gating ring between the metal-free and liganded aSlo1 structures (Hite et al., 2017; Tao et al., 2017): the metal-free gating ring moves ~5 Å away from the TM region. Due to the rotational flexibility of the transmembrane (TM) domain as observed in metal-free aSlo1 (Hite et al., 2017), the TM helices will be displaced proportionally to the distance to the ion-conducting pore (e.g. 6 Å at helix S1 and 9 Å at helix S0) (Figure S5A). As a result, only helices S5, S6 and pore domain were determined together with the gating ring using the C2 symmetry.

**Figure 2:**
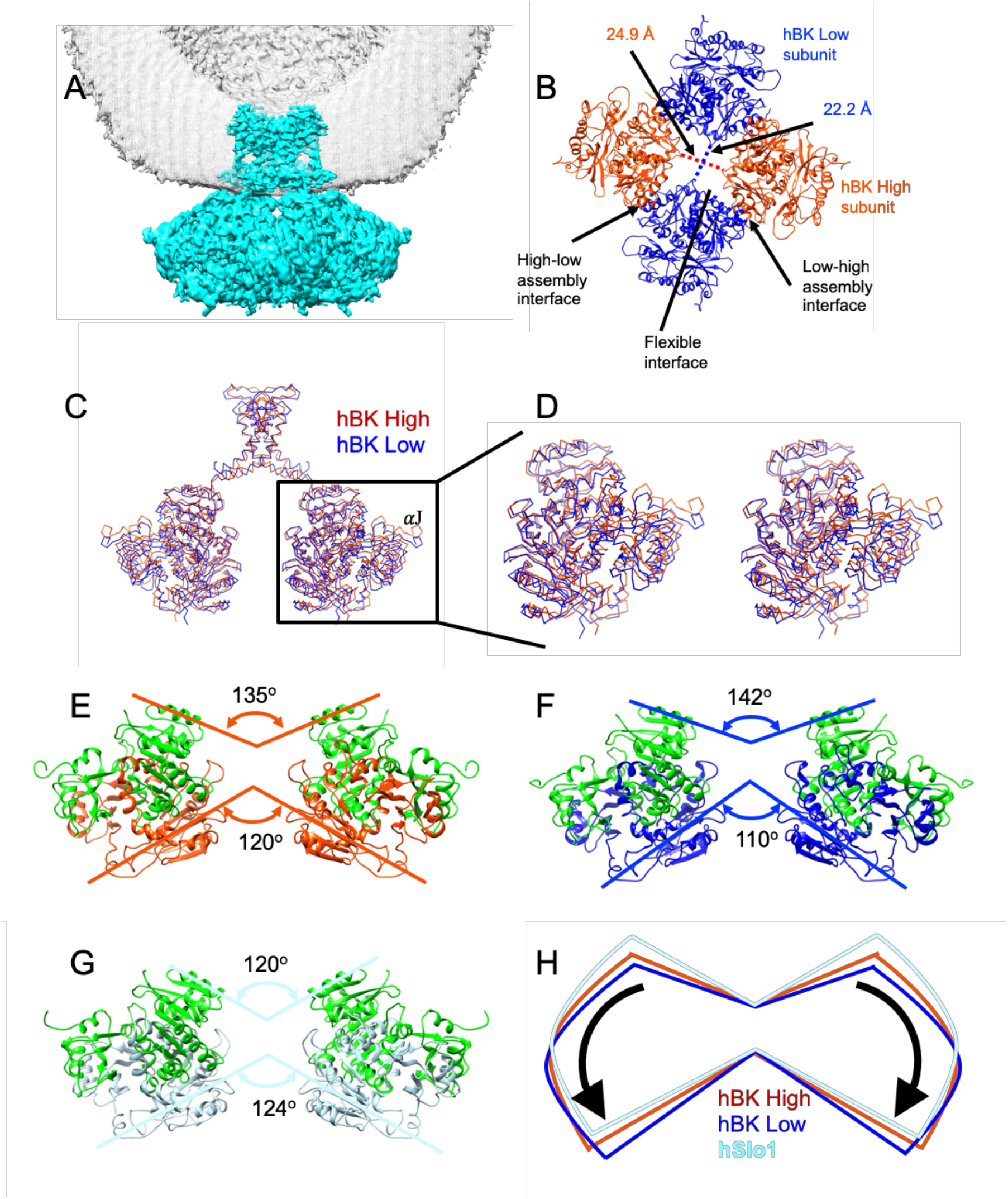
The structure of hBK in liposomes. (A) hBK cryo-EM density map. Lipid membrane is shown in gray mesh. For clarity, only half of the lipid membrane is shown. (B) View of the gating ring in hBK from the extracellular side. The diagonal distances between the C_α_ atoms of Val 785 are indicated. (C) Superposition of hBK High (orange red) and Low (blue) subunits. (D) Stereo view of the superimposed hBK High and Low subunits. (E-G) Side views of hBK High (E), hBK Low (F) and hSlo1-GR (G) (light blue) (PDB: 3NAF). To distinguish RCK1 and RCK2 domains, the RCK1 domains in hBK were colored green. The RCK2 domains in hBK high subunits are aligned to those in hSlo1-GR. (H) Cartoon to show the relative rotation among hBK High and Low, and hSlo1-GR. Related to Figures S2-4, 6, Movies S1-5.

Compared with the structure of the isolated hSlo1 gating ring (hSlo1-GR) (Wu et al., 2010), a significant conformational change was observed (Figure 2 E-H, Movie S2). In general, the RCK1 domains in hBK open like the petals of a flower, but the extent of opening is different in hBK High and Low subunits. The two RCK1 domains in hBK High subunits opens 15 degrees more than that in hSlo1-GR at the top (close to the membrane), while the RCK1 domains opens 22 degrees more than that in hSlo1-GR (Figure 2E-G). As RCK1 domains open, the RCK2 domains also undergo conformational changes. The RCK2 domains at the bottom (away from the membrane) in hBK High subunits close by 4 degrees more than the corresponding domains of hSlo1-GR, while the RCK2 domains of hBK close 14 degrees more than that of hSlo1-GR (Figure 2E-G). It is worth noting that the extent of opening of RCK1 at the top differs from the extent of closing of RCK2 at the bottom even in the same subunit. This difference is not due to the relative movement between RCK1 and RCK2 domains in each subunit. As shown in Figure 3 A, the flexible interfaces formed by the RCK1 and RCK2 domains in hBK High and Low subunits are the same after alignment. The difference of the opening at the top and closing at the bottom in hBK High and Low subunits is due to the rotation and tilting of each subunit as shown in Movie S3. As a result, the High-low and Low-High assembly interfaces formed between neighboring subunits remains the same. The interacting residues (I441, M442, I445, H468, L880, L883, I879 and F948) aligned well with each other (Figure 3C). In short the flexible and assembly interfaces in hBK remains the same in hBK High and Low subunits. As hBK channels were reconstituted into liposomes while isolated hSlo1 gating ring formed crystals, some changes were observed in both the flexible and assembly interfaces (Figure 3 B&D). At the flexible interfaces, the positions of the helix-crossover domain (aF-turn-aG) that interlocks the two RCK domains changes from hSlo1-GR to hBK, but the folding is maintained. At the assembly interfaces, the aD and aE helices in hBK Low subunits shifts down (away from the membrane) with respect to those in hSlo1-GR (Figure 3D and Movie S4). The changes were not significant, thus the interaction between neighboring subunits shouldn’t change.

**Figure 3:**
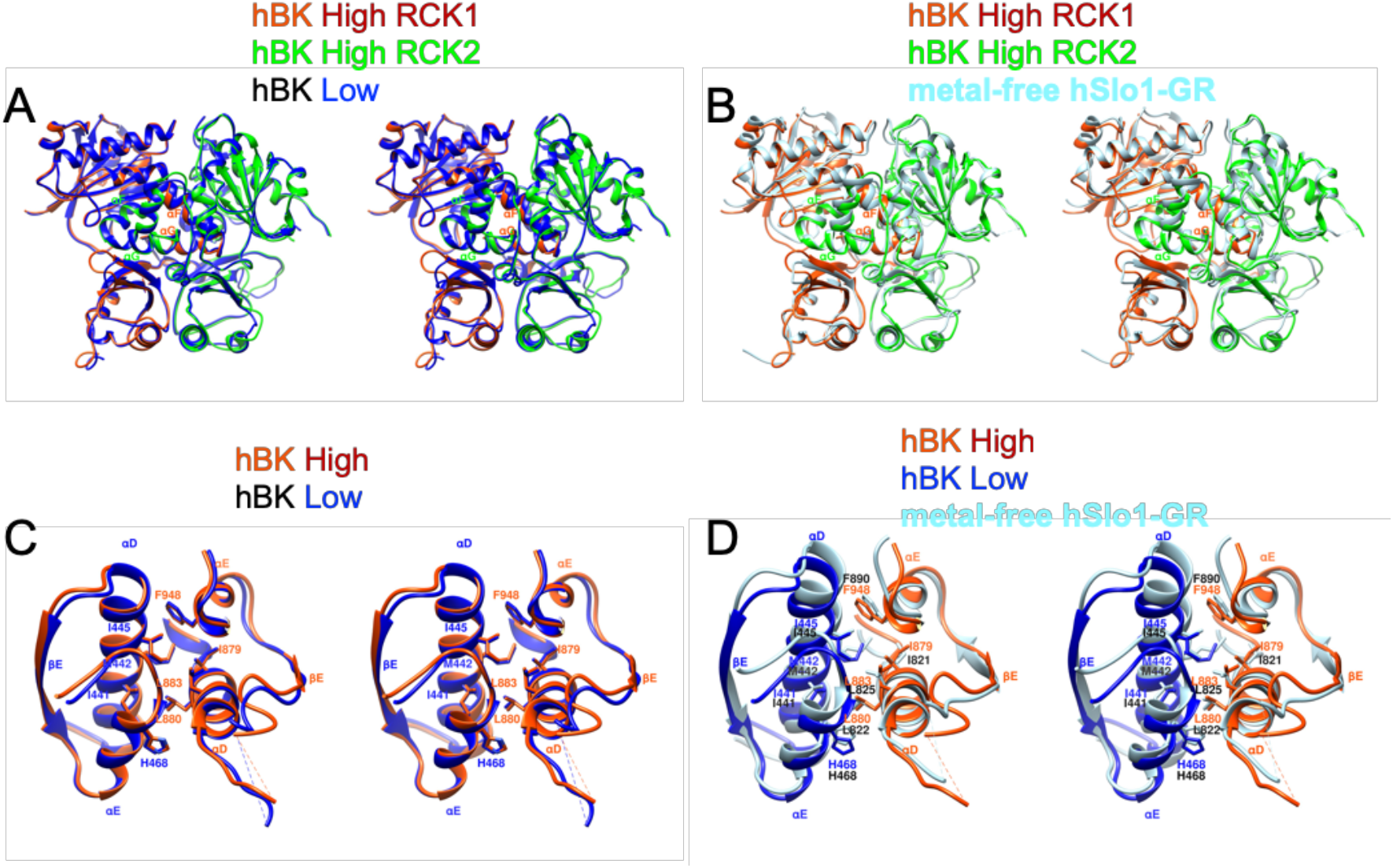
Stereo views of assembly and flexible interfaces in hBK. (A) Flexible interfaces in hBK. hBK High (orange red) and Low (blue) subunits are superimposed. To show the flexible interface, RCK2 in hBK High subunit is colored green. These views are similar to that shown in Figure 2B except the subunit was rotated slightly to reduce the overlap of RCK1 and RCK2. (B) Same as A except the replacement of hBK Low with hSlo1-GR (light blue, PDB: 3NAF) subunit. (C) Assembly interface in hBK as defined in Figure 2B. hBK High (orange red) and Low (blue) subunits are superimposed. (D) Same as C except the replacement of hBK Low with hSlo1-GR (light blue, PDB: 3NAF) subunit. The αD and αE helices in RCK2 of hSlo1-GR were aligned to those of hBK.

### An intermediate state of Ca^2+^ sensing

There exist two Ca^2+^-binding sites in hBK: the Ca^2+^ bowl and the RCK1 Ca^2+^-binding site. At the Ca^2+^ bowl, residues D953, D955 and N449 (D905, D907 and N438 in aSlo1) and backbone carbonyl of D950 and Q947 (D902 and Q899 and in aSlo1) are expected to provide the basis to coordinate a Ca^2+^ ion as seen in liganded aSlo1 (Figure 4B and Supplementary Figure S6E). In the absence of Ca^2+^, the Ca^2+^ bowl in hBK (same conformation in both hBK High and Low subunits) differs significantly from that in hSlo1-GR (Figure 4 A): the N449 and D953 moved toward the Ca^2+^ binding pocket, while D955 swings away from the Ca^2+^ binding pocket. Similar movement of those three residues (N438, D905 and D907) were observed in metal-free aSlo1 (Figure 4B). At the low-affinity Ca^2+^ binding site, residues D367, E535, S600 and S533 (D356, E525, and E591 in aSlo1) (S533 in hBK replaces G523 in aSlo1) and the backbone carbonyl of R503 (R503 and G523 in aSlo1) are expected to provide coordination of a Ca^2+^ ion as seen in aSlo1 (Figure S6F). In metal-free hBK, the D367 moved closer to the Ca^2+^ binding pocket (Figure 4C) than the D367 in hSlo1-GR. As seen at the Ca^2+^ bowl, similar movement of D367 (D356 in aSlo1) was observed in metal-free aSlo1 (Figure 4D). In short, both Ca^2+^ binding sites in hBK differ from those in hSlo1-GR, but agree with those in aSlo1.

**Figure 4:**
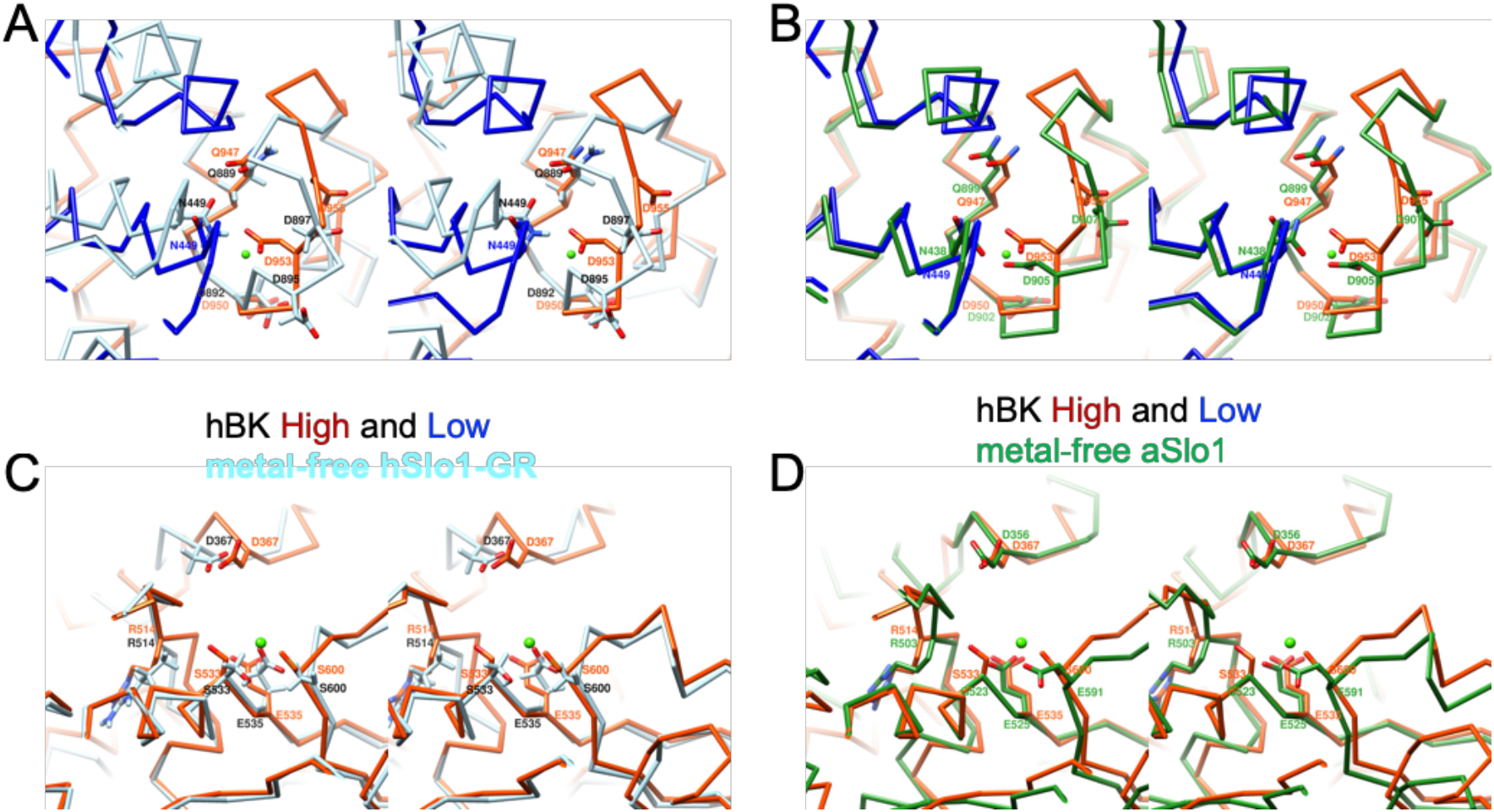
Stereo views of Ca^2+^-binding sites in hBK. (A) The Ca^2+^ bowl in metal-free hBK (orange red and blue) and metal-free hSlo1-GR (light blue). (B) Same as A except metal-free hSlo1-GR was replaced with metal-free aSlo1 (green). (C) The RCK1 Ca^2+^-binding site in metal-free hBK and metal-free hSlo1-GR. (D) Same as C except metal-free hSlo1-GR was replaced with metal-free aSlo1 (green). Ca^2+^ coordinating residues are labeled, and the position of Ca^2+^ ion in liganded aSlo1 was represented by a green sphere. Related to Figure S6.

### Rotational flexibility in the TM region

In the analysis of metal-free aSlo1 cryo-EM data, alternative 3D maps were obtained that show an 8-degree variation in rotation angle of the TM region relative to the gating ring. We obtained only one 3D class, but presumably due to a similar rotational flexibility of the transmembrane (TM) domain, helices S0-S4 cannot be well resolved in the EM density map with a C2 symmetry. Applying a C4 symmetry however produced a stronger signal. Our hBK structure shows a total rotation of 12 degrees relative to the major 3D class from aSlo1 (Figure 5A). The helix S0 observed in metal-free aSlo1 is not visible in hBK as it is the farthest from the symmetry axis. As shown in Figure 5A, the VSD is more extended compare with that in metal-free aSlo1. As metal-free aSlo1 was solvated in detergents, the VSDs may adopt a more compact conformation. In this study, hBK was reconstituted into liposomes, the lipid bilayer environment was restored. As the lipid membrane is fluidic, the protein has more flexibility, thus the VSDs adopt a more extended conformation.

**Figure 5:**
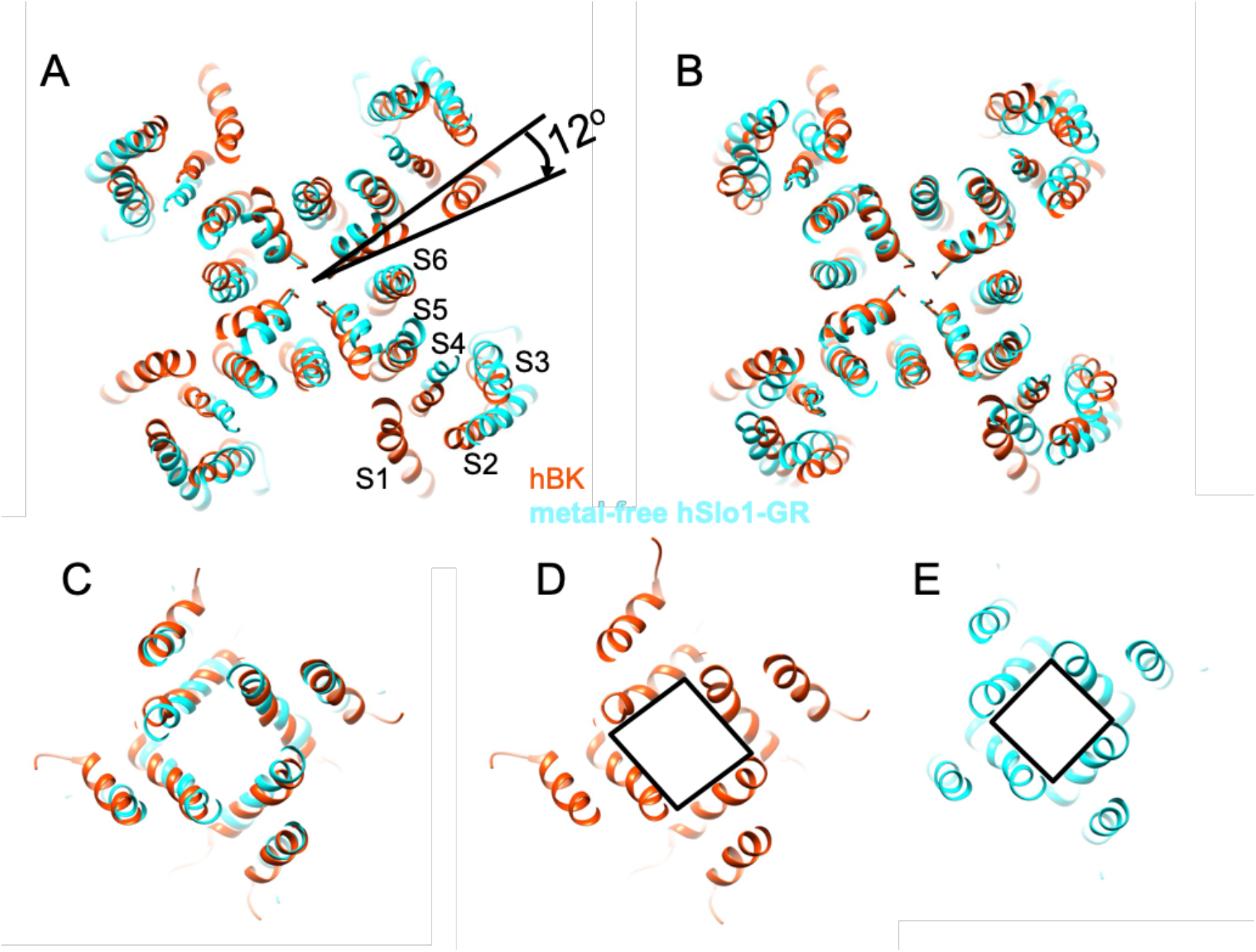
Comparison of the TM region in metal-free hBK and metal-free aSlo1. (A) The TM region in hBK (orange red) is rotated by 12 degrees clockwise with respect to the main 3D class from the aSlo1 dataset (cyan). (B) aSlo1 is shown rotated by 12 degrees to overlay with hBK. All views are from the extracellular side. (C) Superposition of the intracellular helix bundle crossing region of hBK and aSlo1. (D-E) The intracellular helix bundle crossing region of hBK and aSlo1, respectively. Related to Figure S5.

Upon rotating the aSlo1 TM region by 12 degrees, the pore domain of aSlo1 overlapped with that of hBK (Figure 5B). As this pore domain is closer to the central symmetry axis of the channel and experience less blurring (Figure S5), the intensity is much stronger that the intensity in the VSD regions (Figure S5B-C). Thus, the pore domain is visible in the reconstruction with a C2 symmetry (Figure 2 & 5C-D). The intracellular helix bundle crossing of hBK forms a parallelogram instead of a square in aSlo1 (Figure 5 C-E). This is due to the movement of the RCK domains in the gating ring region (Figure 2C-D).

## DISCUSSION

Each BK channel α-subunit contains two RCK domains in its large intracellular C-terminal region. The X-ray crystallographic structure of metal-free hSlo1-GR reveals a gating ring formed by 4 RCK1 and 4 RCK2 domains (Wu et al., 2010) similar to those observed in the *E. coli* K^+^ channel (Jiang et al., 2001), the MthK channel (Jiang et al., 2002a, b; Ye et al., 2006) and aSlo1 channel (Hite et al., 2017; Tao et al., 2017). When hBK channels were reconstituted into liposomes in the absence of Ca^2+^, the four-fold symmetry was broken and a two-fold symmetry was observed. In each hBK subunit, the fold is the same as that in hSlo1-GR (Figure 3B). To accommodate the broken symmetry, the assembly interfaces between neighboring subunits twisted as shown in Figure 3D providing the interactions between critical residues were not changed. As for the Ca^2+^-binding sites, larger conformation changes were observed between hBK and hSlo1-GR. At the Ca^2+^ bowl, residue N449 and D953 moved toward the Ca^2+^ binding pocket and D955 swung away from the Ca^2+^ binding pocket. At the RCK1 Ca^2+^ binding site, D367 moved toward the Ca^2+^ binding pocket. The origin of the difference may be due to crystal constraints as both Ca^2+^ binding sites in hBK are similar to those in aSlo1 determined using cryo-EM (Figure 3). As the gating ring in aSlo1 also has a four-fold symmetry, a similar opening of RCK1 and closing of RCK2 was observed from the metal free aSlo1 to hBK (Movie S5, Figure S6A-D). The hBK High subunits have a similar distance to the membrane as those in aSlo1, while the hBK Low subunits are further away from the membrane (Figure S6). The origin of the down movement of the hBK Low subunits or the two-fold symmetry instead of the four-fold symmetry might lie in the restoration of the lipid membrane environment.

Based on the metal-free and liganded aSlo1 structures, Hite *et al* proposed a model of Slo1 gating by Ca^2+^ and voltage(Hite et al., 2017). Here, the model was modified to include the hBK structure (Figure 6). In the presence of Ca^2+^ under positive transmembrane potential (e.g. 120-240 mV), the gating ring adopts a compact conformation, similar to liganded aSlo1 in detergents (Tao et al., 2017). The VSDs are up and the channel is open. In the absence of Ca^2+^, the gating ring adopts a loose conformation and is away from the membrane, as seen in ligand-free aSlo1 in detergents (Hite et al., 2017). As the transmembrane potential drops to zero, two VSDs move down (maybe not fully down as in the hyperpolarized membrane), thus push the two opposing subunits away from the membrane (hBK Low subunits). This is consistent with that the opening step has been shown to occur cooperatively among the subunits(Mannuzzu and Isacoff, 2000; Pathak et al., 2005) and comprises at least two transitions(Kalstrup and Blunck, 2013). To confirm this hypothesis, at least the structure of hBK reconstituted in liposomes in the presence of Ca^2+^ is needed. Of course, the structures with VSDs in the down conformation are also required to obtain a real understanding of the coupling between Ca^2+^ sensing, voltage sensing and channel opening. A negative potential will be applied to trap hBK channels in a fully closed states with and without Ca^2+^ in the near future using the RSC method.

**Figure 6:**
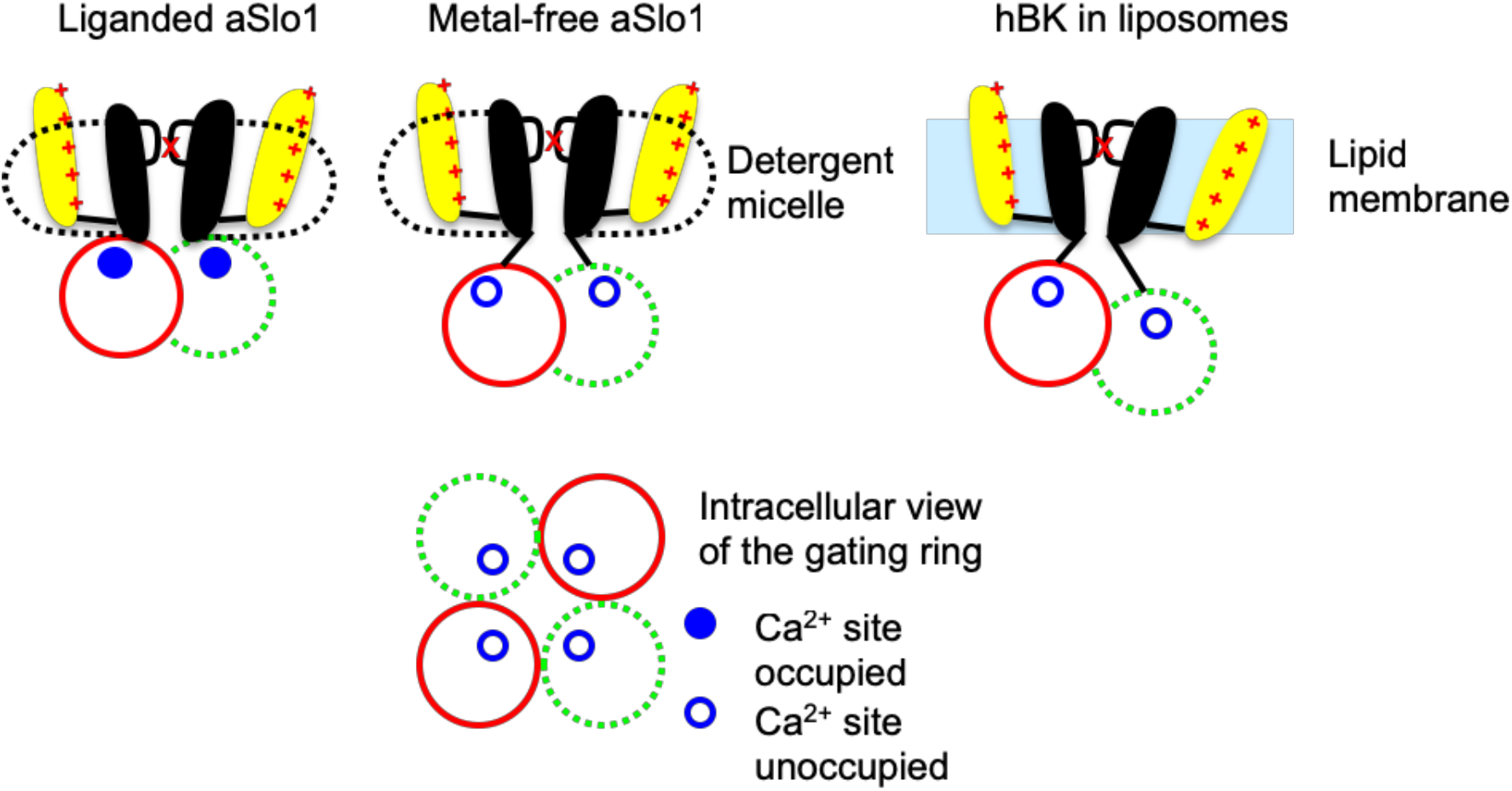
Comparison of metal-free hBK, metal-free aSlo1 and liganded aSlo1. The model proposed by Hite et al (Hite et al., 2017)was modified to include the hBK structure. In the presence of Ca^2+^ under positive transmembrane potential (e.g. 240 mV), the gating ring adopts a compact conformation and the VSDs are expected to be up, similar to the liganded aSlo1 in detergents. When there is no Ca^2+^, the gating ring region adopts a loose conformation as seen ligand-free aSlo1 in detergents. As the transmembrane potential drops to zero, two VSDs move downward resulted in the downward movement of two subunits in the gating ring as seen in hBK. Two adjacent subunits instead of two opposing subunits are shown in the cartoon.

The proteoliposomes used for structure determination ranged from 15 to 70 nm in diameter with the mean diameter at 20 nm. The question therefore arises, whether the membrane curvature in liposomes of different sizes would affect the observed hBK structure, especially the C2 symmetry. To address this question, three hBK structures were reconstructed from small (15-19 nm), intermediate (19-25 nm) and large (25-70 nm) proteoliposomes respectively (Figure S3). The gating ring agrees well in the three reconstructions, and all show the C2 symmetry. A parallel reconstruction with the liposome information unsubtracted particle images, illustrates the curved membrane (Figure S3). For examination of the TM region, C4 symmetry was applied as discussed before. The reconstructions from hBK reconstituted in intermediate and large liposomes show similar VSD helices in addition to the pore region as that in the reconstructions from hBK reconstituted in all the liposomes. The pore region in the reconstruction from hBK reconstituted from small liposomes agrees well with other reconstructions, but the VSD is much noisier. Thus, the membrane curvature only affects the VSD region when the liposomes are smaller than 19 nm in diameter.

## METHOD DETAILS

### Expression, purification and reconstitution of human BK proteins

The BK proteins were expressed and purified as previously described (Wang and Sigworth, 2009). Briefly, full-length human Slo protein (GI:507922) carrying an N-terminal FLAG tag was stably expressed in HEK293 cells. The BK protein was purified using an anti-FLAG affinity column (Sigma Aldrich), where the detergent dodecyl maltoside (DDM) (Anatrace) was exchanged with decyl maltoside (DM) (Anatrace) before elution with FLAG peptide (Sigma Aldrich). Size-exclusion chromatography (SEC) on a Superose 6 column was used to further purify the BK channel.

To reconstitute the BK protein for structural study, the major SEC peak (Figure S1) was mixed with DM-solubilized POPC (1-palmitoyl-2-oleoyl-sn-glycero-3-phosphocholine) lipid (Avanti Polar Lipids, Inc.) (DM: POPC = 3:1) giving a final protein-to-lipid molar ratio of 1:1,000. Gel filtration on a hand-packed 24ml (10mm I.D and 300mm height) was used to remove detergent with running buffer containing 20 mM HEPES, pH 7.3, 150 mM KCl, 2 mM EDTA. To reconstitute the BK for functional assay, the purified BK were reconstituted into liposomes as previously described (Li et al., 2015). Briefly, the POPC and POPG (1-palmitoyl-2-oleoyl-sn-glycero-3-phospho-(1’-rac-glycerol)) lipid mixture (3:1 molar ratio) in chloroform was dried under nitrogen for 30 min, and rehydrated and sonicated in Fisher Scientific FS20H Ultrasonic Cleaner (Fisher Scientific) in buffer A (20 mM HEPES, pH 7.3, 150 mM KCl) to a final lipid concentration of 10 mM. The resulting solution was mixed with detergent DM (lipid: DM = 1: 1) for 30 min then mixed with the purified BK channels to a final protein concentration of 0.05 mg/ml. The detergent was removed by dialysis in 12-14 KD cut-off dialysis membrane (Spectrum Labs) against the buffer A at 4 °C for 3 days, with daily buffer exchange. Empty liposomes were prepared in the same manner without the addition of protein prior to dialysis. The final liposome prep was concentrated to ~2 mM total lipid concentration.

### Flux assay to assess the function of reconstituted BK proteins and estimate the transmembrane potential

Empty liposomes or proteoliposomes were diluted 100-fold into a buffer containing 150 mM NaCl, 20 mM HEPES pH 7.3, or a buffer containing 136.5 mM NaCl, 13.5 mM KCl, 20 mM HEPES, pH 7.3. ACMA (9-amino-2-methoxy-6-chloroacridine, Sigma-Aldrich) stock was added to a final concentration of 2 μM. This yielded a final K^+^ concentration of 1.5 mM or 15 mM outside liposomes. Fluorescence intensity was measured every 1 s using Fluorolog^®^-3 Spectroflorometer (HORIBA Scientific) with excitation at 395 nm and emission at 490 nm. The protonophore carbonyl cyanide m-chlorophenyl hydrazone (CCCP, Sigma-Aldrich) was added to a final concentration of 1 μM. Valinomycin (Sigma-Aldrich) was added to a final concentration of 20 nM.

### Cryo-EM sample preparation and image collection

To obtain highly spherical liposomes, we swelled proteoliposomes by repeated osmotic shocks, adding water to the liposome suspension (11%, 14%, 18%, and 24% of the original volume) at 1 h intervals at 4°C. Then 2μl of BK proteoliposomes at a concentration of 2 mM lipid and 1mg/ml protein was applied onto a glow-discharged perforated carbon grids (CFlat R2/2, EMS), and incubated for 5-10 min at 22 °C and 100% relative humidity in an FEI Vitrobot Mark IV (FEI). After incubation, the sample was blotted from the side and 2μl of buffer containing 10 mM HEPES, pH 7.3, 75 mM KCl, and 1 mM EDTA was applied onto the TEM grid to swell proteoliposomes. Then grids were blotted for 7 s with force 0 and fast frozen in liquid ethane cooled by liquid nitrogen. Grids were then transferred to an FEI Titan Krios electron microscope operating at an acceleration voltage of 300 keV with an energy filter (20eV slit width). Images were recorded in an automated fashion on a Gatan K2 Summit (Gatan) detector set to super-resolution mode with a super-resolution pixel size of 0.525 Å using Leginon (Suloway et al., 2005). The dose rate on the camera was set to be ~8 counts per physical pixel per second. The total exposure time was 8.6 s (0.2 s/frame), leading to a total accumulated dose of 60 electrons per Å^2^ on the specimen.

### Image processing and map calculation

Image processing is illustrated in Figure S2A. First, Dose-fractionated super-resolution image stacks (0.525 Å/pixel) were motion-corrected with MotionCor2 (Zheng et al., 2016). Each frame in the image stack was divided into 5×5 patches for anisotropic image motion correction and dose weighting was carried out to calculate the motion-corrected image. Then the parameters of the contrast transfer function were estimated by CTFFIND4 (Rohou and Grigorieff, 2015). Following motion correction, the liposome contribution was removed with a model based on the average POPC membrane profile from 280 micrographs (The model was obtained using the Hankel transform as previously described (Wang et al., 2006)). Then, approximately 4,800 particles from 350 micrographs were interactively selected and classified using RELION(Scheres, 2012). Ten representative classes were selected as the templates for automated particle picking using RELION. The automated picked particles were screened using homemade MATLAB (MathWorks) programs to exclude particles more than 80 Å away from any liposome, which resulted in 275,295 particles from 2,786 micrographs. Then 2D and 3D classifications were carried out to select 122,456 particles for cryo-EM map reconstruction. The metal-free aSlo1 was used as the initial model. The EM density map determined with a C1 symmetry clearly showed a two-fold symmetry (Figure S2). With a C2 symmetry imposed, the resolution of the EM density map was 3.61 Å. Then the structure was further refined in CryoSparc (Punjani et al., 2017) to 3.48 Å.

During the liposome-density subtraction step, modeled images of intact liposomes were subtracted from micrographs. In the transmembrane (TM) region, therefore, only the difference between hBK and lipid bilayer density remained, a signal that is much weaker than the true protein density. Thus the signal in the TM region was amplified by a factor of 2.5-4.0 to make the helices in TM region and in the gating ring have similar intensity. To show the position of the membrane, an alternative 3D reconstruction was obtained from a matching particle stack extracted from non-liposome-subtracted micrographs.

The initial model of hBK was predicted using SWISS-MODEL(Waterhouse et al., 2018). Then the structure was docked into the EM map, and flexible loops were removed manually using UCSF Chimera(Pettersen et al., 2004). Then a mask extending by 3 pixels and with a soft edge of 8 pixels was created in RELION, and applied to the EM map. The masked EM map was using for flexible fitting using MDFF (Trabuco et al., 2009) with a C2 symmetry. The refined structure was further refined in real space using phenix.real_space_refine(Adams et al., 2011). Geometric and secondary structure restraints were tightly maintained throughout refinement to minimize over fitting.

## DATA AND SOFTWARE AVAILABILITY

The 3D cryo-EM density maps of hBK and atomic model will be deposited in the Electron Microscopy Data Bank.

## Supporting information

## ACKNOWLEDGMENTS

We thank Drs. William N. Zagotta and Galen Flynn (University of Washington) for the use of the fluorometer, Drs. Hong Z. Zhou, Yanxiang Cui, Wong Hoi Hui, and Ivo Atanasov (UCLA) for the use of the Titan Krios at UCLA, Drs. Eric Gouaux, Claudia S. López, and Craig Yoshioka (Oregon Health & Science University) for the use of the Titan Krios at Oregon Health & Science University, and Dr. Fred J. Sigworth for discussions. This work was supported by the National Institutes of Health [grant number: R01GM096458].

## AUTHOR CONTRIBUTIONS

Conceptualization, L.W.; Methodology, T.L. and L.W.; Software, L.W.; Writing, L.W.; Editing, L. T. and L.W.; Visualization, L. T. and L.W.; Project Administration, L.W.; Funding Acquisition, L.W.

## DECLARATION OF INTERESTS

The authors declare no competing interests.

## References

Adams, P.D., Afonine, P.V., Bunkóczi, G., Chen, V.B., Echols, N., Headd, J.J., Hung, L.W., Jain, S., Kapral, G.J., Grosse Kunstleve, R.W., et al. (2011). The Phenix software for automated determination of macromolecular structures. Methods 55, 94–106.

Autzen, H.E., Myasnikov, A.G., Campbell, M.G., Asarnow, D., Julius, D., and Cheng, Y. (2018). Structure of the human TRPM4 ion channel in a lipid nanodisc. Science (New York, NY) 359,228–232.

Bayburt, T.H., Grinkova, Y.V., and Sligar, S.G. (2002). Self-Assembly of Discoidal Phospholipid Bilayer Nanoparticles with Membrane Scaffold Proteins. Nano Lett 2, 853–856.

Brohawn, S.G., del Marmol, J., and MacKinnon, R. (2012). Crystal structure of the human K2P TRAAK, a lipid- and mechano-sensitive K+ ion channel. Science 335, 436–441.

Chen, Q., She, J., Zeng, W., Guo, J., Xu, H., Bai, X.-C., and Jiang, Y. (2017). Structure of mammalian endolysosomal TRPML1 channel in nanodiscs. Nature 550, 415–418.

Cui, J., Yang, H., and Lee, U.S. (2009). Molecular mechanisms of BK channel activation. Cellular and Molecular Life Sciences (CMLS) 66, 852–875.

Dang, S., Feng, S., Tien, J., Peters, C.J., Bulkley, D., Lolicato, M., Zhao, J., Zuberbühler, K., Ye, W., Qi, L., et al. (2017). Cryo-EM structures of the TMEM16A calcium-activated chloride channel. Nature 552, 426–429.

Gao, Y., Cao, E., Julius, D., and Cheng, Y. (2016). TRPV1 structures in nanodiscs reveal mechanisms of ligand and lipid action. Nature 534, 347–351.

Gonen, T., Cheng, Y., Sliz, P., Hiroaki, Y., Fujiyoshi, Y., Harrison, S.C., and Walz, T. (2005). Lipid-protein interactions in double-layered two-dimensional AQP0 crystals. Nature 438, 633–638.

Gribkoff, V.K., Starrett, J.E., and Dworetzky, S.I. (2001). Maxi-K potassium channels: Form, function, and modulation of a class of endogenous regulators of intracellular calcium. Neuroscientist 7, 166–177.

Hilgemann, D.W. (2003). Getting ready for the decade of the lipids. Annu Rev Physiol 65, 697–700.

Hille, B., Dickson, E.J., Kruse, M., Vivas, O., and Suh, B.-C. (2015). Phosphoinositides regulate ion channels. Biochim Biophys Acta 1851, 844–856.

Hite, R.K., Tao, X., and MacKinnon, R. (2017). Structural basis for gating the high-conductance Ca^2+^-activated K^+^ channel. Nature 541, 52–57.

Horrigan, F.T., and Aldrich, R.W. (2002). Coupling between voltage sensor activation, Ca^2+^ binding and channel opening in large conductance (BK) potassium channels. J Gen Physiol 120, 267–305.

Jackson, S.M., Manolaridis, I., Kowal, J., Zechner, M., Taylor, N.M.I., Bause, M., Bauer, S., Bartholomaeus, R., Bernhardt, G., Koenig, B., et al. (2018). Structural basis of small-molecule inhibition of human multidrug transporter ABCG2. Nat Struct Mol Biol 25, 333–340.

Jensen, K.H., Brandt, S.S., Shigematsu, H., and Sigworth, F.J. (2016). Statistical modeling and removal of lipid membrane projections for cryo-EM structure determination of reconstituted membrane proteins. J Struct Biol 194, 49–60.

Jiang, Y., Pico, A., Cadene, M., Chait, B.T., and MacKinnon, R. (2001). Structure of the RCK domain from the E. coli K^+^ channel and demonstration of its presence in the human BK channel. Neuron 29, 593–601.

Jiang, Y.X., Lee, A., Chen, J.Y., Cadene, M., Chait, B.T., and MacKinnon, R. (2002a). Crystal structure and mechanism of a calcium-gated potassium channel. Nature 417, 515–522.

Jiang, Y.X., Lee, A., Chen, J.Y., Cadene, M., Chait, B.T., and MacKinnon, R. (2002b). The open pore conformation of potassium channels. Nature 417, 523–526.

Kalstrup, T., and Blunck, R. (2013). Dynamics of internal pore opening in K_v_, channels probed by a fluorescent unnatural amino acid. Proceedings of the National Academy of Sciences 110, 8272–8277.

Katayama, H., Wang, J., Tama, F., Chollet, L., Gogol, E.P., Collier, R.J., and Fisher, M.T. (2010). Three-dimensional structure of the anthrax toxin pore inserted into lipid nanodiscs and lipid vesicles. Proc Natl Acad Sci U S A 107, 3453–3457.

Lee, A.G. (2011). Biological membranes: the importance of molecular detail. Trends Biochem Sci 36, 493–500.

Li, M., Tonggu, L., Tang, L., and Wang, L. (2015). Effects of N-glycosylation on hyperpolarization-activated cyclic nucleotide-gated (HCN) channels. Biochem J 466, 77–84.

Long, S.B., Tao, X., Campbell, E.B., and MacKinnon, R. (2007). Atomic structure of a voltage-dependent K+ channel in a lipid membrane-like environment. Nature 450, 376–383.

Mannuzzu, L.M., and Isacoff, E.Y. (2000). Independence and Cooperativity in Rearrangements of a Potassium Channel Voltage Sensor Revealed by Single Subunit Fluorescence. The Journal of General Physiology 115, 257–268.

McGoldrick, L.L., Singh, A.K., Saotome, K., Yelshanskaya, M.V., Twomey, E.C., Grassucci, R.A., and Sobolevsky, A.I. (2018). Opening of the human epithelial calcium channel TRPV6. Nature 553, 233–237.

Miller, A.N., and Long, S.B. (2012). Crystal structure of the human two-pore domain potassium channel K2P1. Science 335, 432–436.

Pathak, M., Kurtz, L., Tombola, F., and Isacoff, E. (2005). The Cooperative Voltage Sensor Motion that Gates a Potassium Channel. The Journal of General Physiology 125, 57–69.

Pettersen, E.F., Goddard, T.D., Huang, C.C., Couch, G.S., Greenblatt, D.M., Meng, E.C., and Ferrin, T.E. (2004). UCSF Chimera - A visualization system for exploratory research and analysis. J Comput Chem 25, 1605–1612.

Punjani, A., Rubinstein, J.L., Fleet, D.J., and Brubaker, M.A. (2017). CryoSPARC: Algorithms for rapid unsupervised cryo-EM structure determination. Nat Methods 14, 290–296.

Roh, S.-H., Stam, N.J., Hryc, C.F., Couoh-Cardel, S., Pintilie, G., Chiu, W., and Wilkens, S. (2018). The 3.5-Å CryoEM Structure of Nanodisc-Reconstituted Yeast Vacuolar ATPase Vo Proton Channel. Mol Cell 69, 993–1004.e1003.

Rohou, A., and Grigorieff, N. (2015). CTFFIND4: Fast and accurate defocus estimation from electron micrographs. J Struct Biol 192, 216–221.

Scheres, S.H.W. (2012). RELION: Implementation of a Bayesian approach to cryo-EM structure determination. J Struct Biol 180, 519–530.

Schmidt, D., Jiang, Q.X., and MacKinnon, R. (2006). Phospholipids and the origin of cationic gating charges in voltage sensors. Nature 444, 775–779.

Schreiber, M., and Salkoff, L. (1997). A novel calcium-sensing domain in the BK channel. Biophys J 73, 1355–1363.

Shi, J.Y., Krishnamoorthy, G., Yang, Y.W., Hu, L., Chaturvedi, N., Harilal, D., Qin, J., and Cui, J.M. (2002). Mechanism of magnesium activation of calcium-activated potassium channels. Nature 418, 876–880.

Srivastava, A.P., Luo, M., Zhou, W., Symersky, J., Bai, D., Chambers, M.G., Faraldo-Gómez, J.D., Liao, M., and Mueller, D.M. (2018). High-resolution cryo-EM analysis of the yeast ATP synthase in a lipid membrane. Science (New York, NY) 360.

Su, Z., Brown, E.C., Wang, W., and MacKinnon, R. (2016). Novel cell-free high-throughput screening method for pharmacological tools targeting K+channels. Proc Natl Acad Sci U S A 113, 5748–5753.

Suloway, C., Pulokas, J., Fellmann, D., Cheng, A., Guerra, F., Quispe, J., Stagg, S., Potter, C.S., and Carragher, B. (2005). Automated molecular microscopy: The new Leginon system. J Struct Biol 151, 41–60.

Sweet, T.-B., and Cox, D.H. (2008). Measurements of the BK_Ca_ Channel’s High-Affinity Ca^2+^ Binding Constants: Effects of Membrane Voltage. The Journal of General Physiology 132, 491–505.

Tao, X., Hite, R.K., and MacKinnon, R. (2017). Cryo-EM structure of the open high-conductance Ca^2+^-activated K^+^ channel. Nature 541, 46–51.

Taylor, N.M.I., Manolaridis, I., Jackson, S.M., Kowal, J., Stahlberg, H., and Locher, K.P. (2017). Structure of the human multidrug transporter ABCG2. Nature 546, 504–509.

Trabuco, L.G., Villa, E., Schreiner, E., Harrison, C.B., and Schulten, K. (2009). Molecular dynamics flexible fitting: A practical guide to combine cryo-electron microscopy and X-ray crystallography. Methods 49, 174–180.

Wang, L., Bose, P.S., and Sigworth, F.J. (2006). Using cryo-EM to measure the dipole potential of a lipid membrane. Proc Natl Acad Sci U S A 103, 18528–18533.

Wang, L., and Sigworth, F.J. (2009). Structure of the BK potassium channel in a lipid membrane from electron cryomicroscopy. Nature 461, 292–295.

Waterhouse, A., Bertoni, M., Bienert, S., Studer, G., Tauriello, G., Gumienny, R., Heer, F.T., de Beer, T.A P., Rempfer, C., Bordoli, L., et al. (2018). SWISS-MODEL: homology modelling of protein structures and complexes. Nucleic Acids Res 46, W296–W303.

Wu, Y., Yang, Y., Ye, S., and Jiang, Y. (2010). Structure of the gating ring from the human large-conductance Ca^2+^-gated K^+^ channel. Nature 466, 393–397.

Xia, X.-M., Zeng, X., and Lingle, C.J. (2002). Multiple regulatory sites in large-conductance calcium-activated potassium channels. Nature 418, 880–884.

Ye, S., Li, Y., Chen, L., and Jiang, Y. (2006). Crystal Structures of a Ligand-free MthK Gating Ring: Insights into the Ligand Gating Mechanism of K+ Channels. Cell 126, 1161–1173.

Yuan, P., Leonetti, M.D., Hsiung, Y., and MacKinnon, R. (2012). Open structure of the Ca^2+^ gating ring in the high-conductance Ca^2+^-activated K^+^ channel. Nature 481, 94–97.

Yuan, P., Leonetti, M.D., Pico, A.R., Hsiung, Y., and MacKinnon, R. (2010). Structure of the human BK channel Ca^2+^-activation apparatus at 3.0 Å resolution. Science 329, 182–186.

Zhang, G., Huang, S.-Y., Yang, J., Shi, J., Yang, X., Moller, A., Zou, X., and Cui, J. (2010). Ion sensing in the RCK1 domain of BK channels. Proceedings of the National Academy of Sciences 107, 18700–18705.

Zheng, S., Palovcak, E., Armache, J.-P., Cheng, Y., and Agard, D. (2016). Anisotropic Correction of Beam-induced Motion for Improved Single-particle Electron Cryo-microscopy. bioRxiv.

