## Supplementary material for "Broken symmetry in the human BK channel"

Figure S1

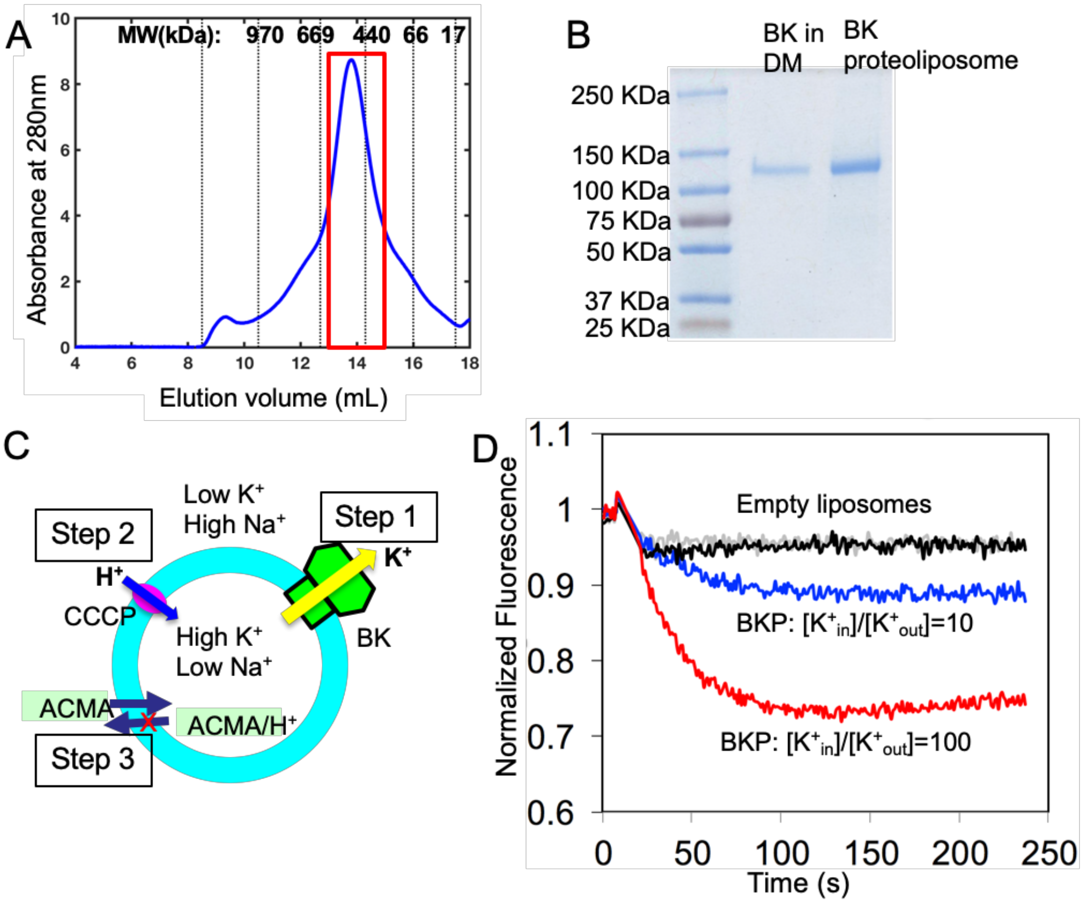


Figure S1. Purification and reconstitution of human BK proteins. (A) FPLC results of the purified hBK protein. Red box marks the fraction collected for structural studies. (B) SDS-PAGE of hBK before and after reconstitution. (C) Cartoon of the system for BK flux assay. Step1. $K^+$ efflux via BK channels. Step2. Proton influx via H+ ionophore CCCP (Carbonyl cyanide m-chlorophenyl hydrazone). Step3. ACMA (Fluorescent dye 9-amion-6-chloro-2-methoxyacridine) is quenched. (D) Fluorescence decreases at different potassium gradients across the lipid bilayer. Related to Figure 1.

Figure S2

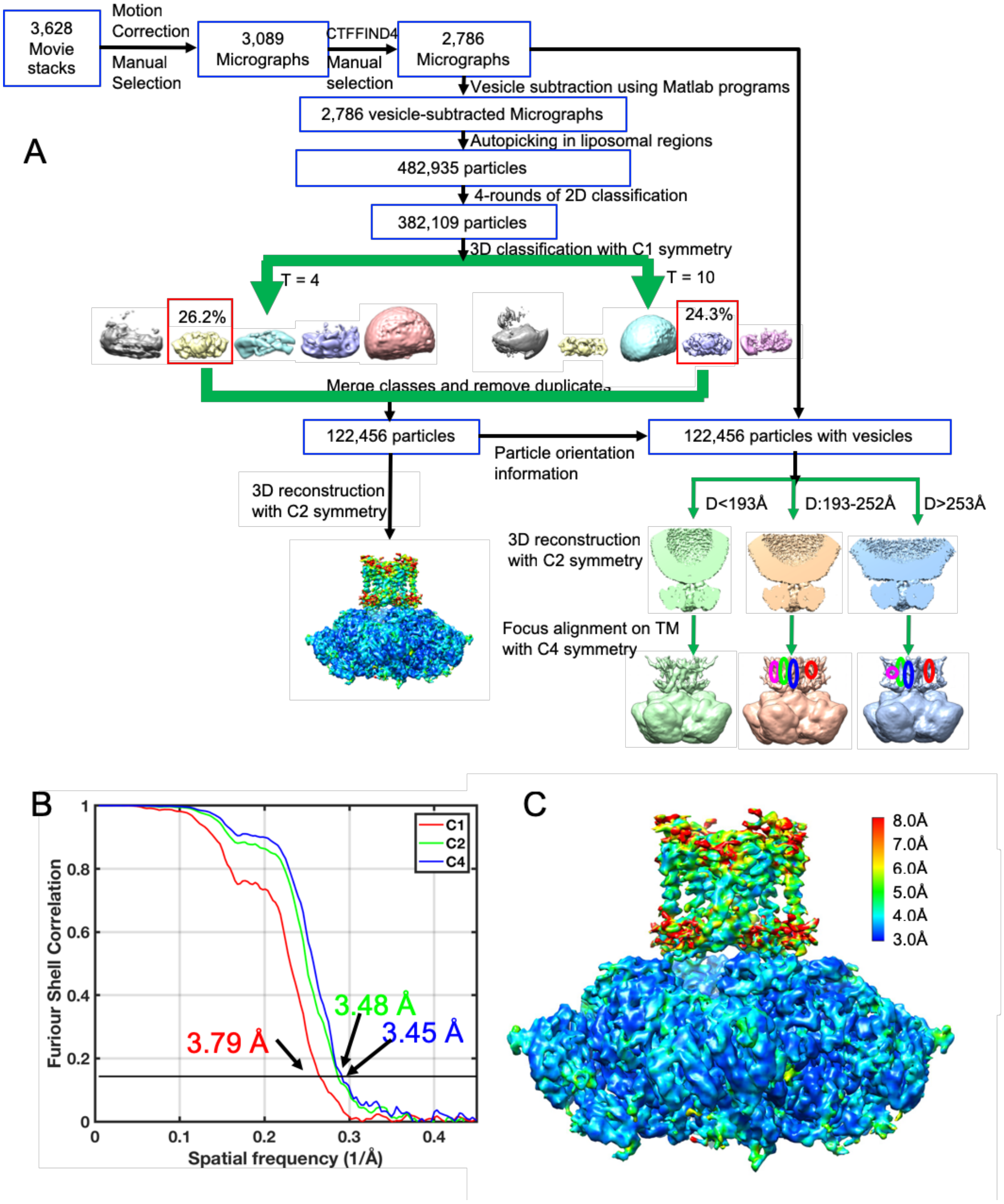


Figure S2. Image processing. (A)Image processing steps. Motion correction of acquired movies (43 frames at 1.4 $e^{-}$/ $Å^2$ /frame) was performed with MotionCor2 (Zheng et al., 2016). Particle selection, classification and reconstruction operations were carried out with Relion (Scheres, 2012). Note that two separate 3D classification operations were performed using values of 4 and

10 for the regularization parameter T. (B) Gold-standard FSC curves from hBK density maps determined with C1, C2 and C4 symmetry. (C) EM density map determined with C2 symmetry colored by local resolution.

Related to Figure 2.

Figure S3

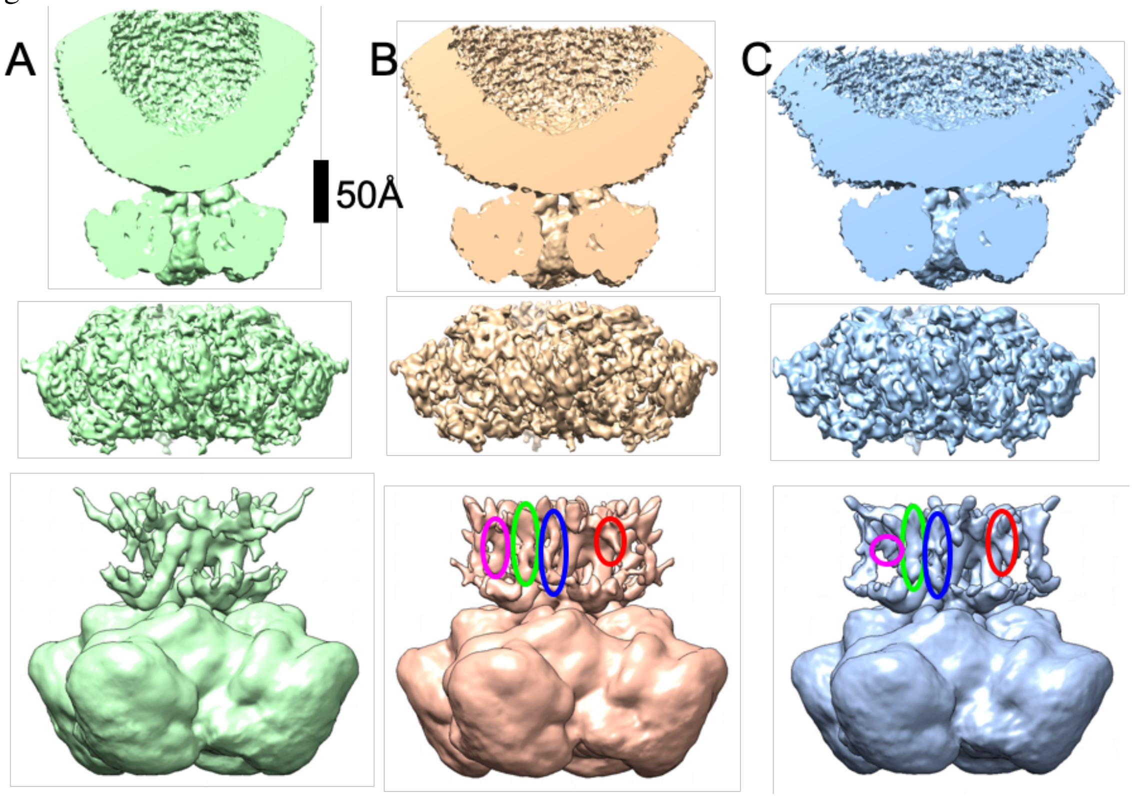


Figure S3. Effect of liposome size on BK structure. (A) Liposome diameter ranges from 150 to 193 Å. (B) Liposome diameter ranges from 193 to 252 Å. (C) Liposome diameter is larger than 252 Å but smaller than 700 Å. Top row: cutaway views of BK in liposomes. Middle row: gating ring of BK. Bottom row: 3D view of BK with TM helices marked. Red, green, blue and magenta ellipses mark helices S1 to S4, respectively. Note: VSD helices are not visible in A.

Related to Figure 2.

Figure S4

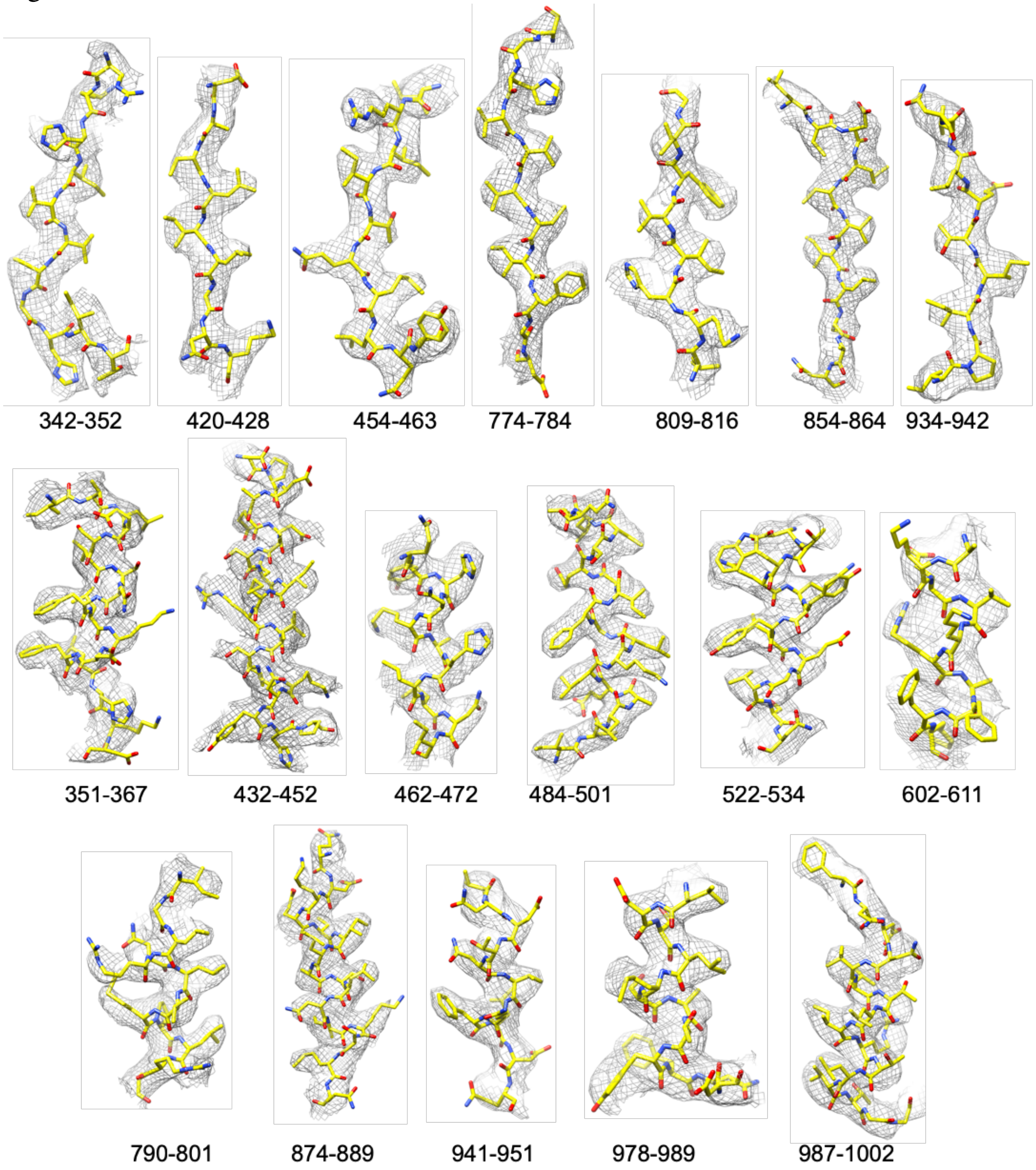


Figure S4. Representative segments of cryo-EM density. Numbers are the residue numbers in hBK.

Related to Figure 2.

Figure S5

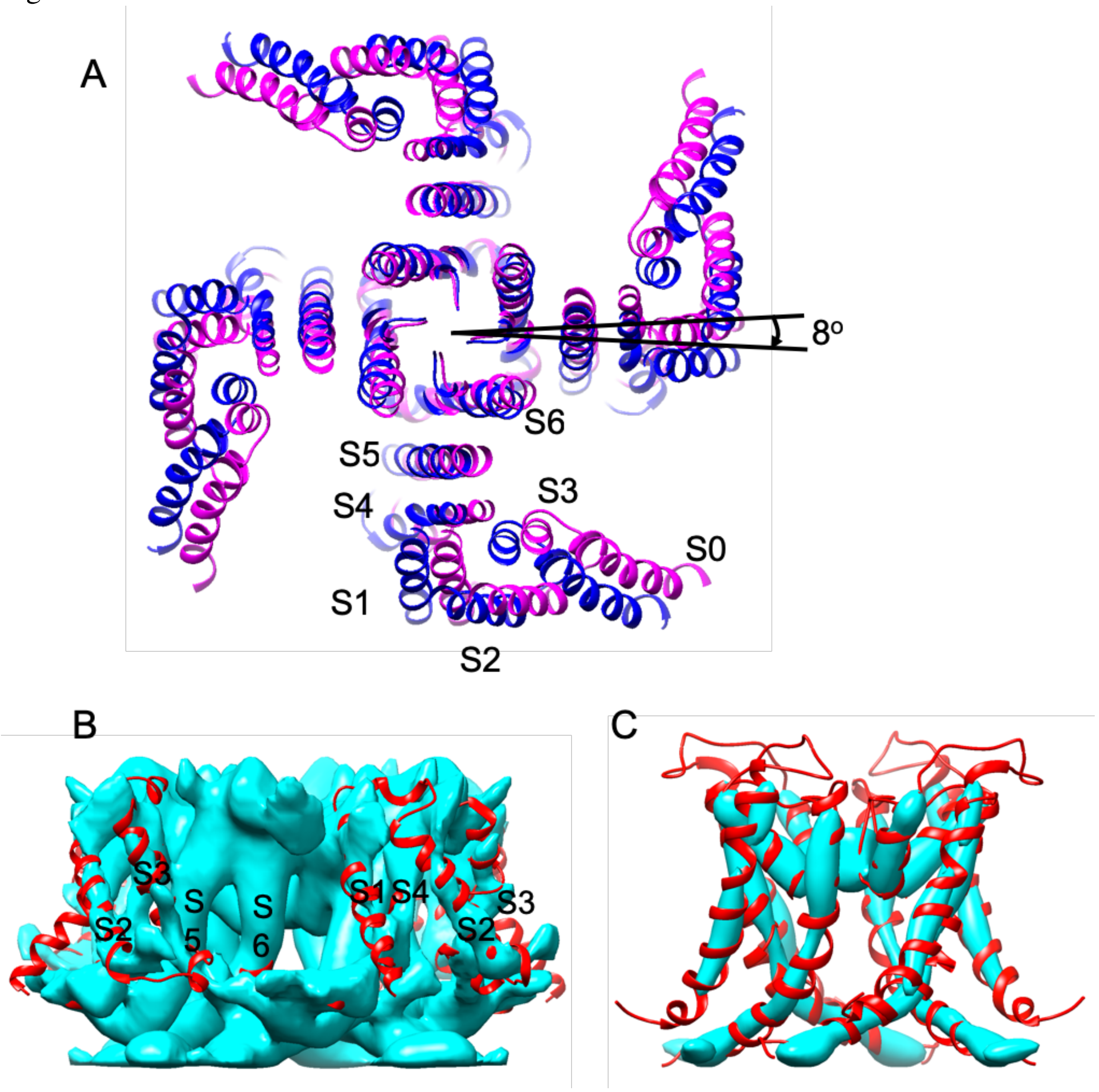


Figure S5. TM region of hBK. (A) Displacement of TM helices in metal-free aSlo1 between the original structure (magenta) and the structure rotated by 8 degree (blue) with respect to the gating ring. Helices S0, S2 and S1 are displaced by 9, 6, and 4 Å, respectively. (B) TM region of hBK at low contour level. (C) The pore domain of hBK. The EM density map was filtered to 6 Å.

Related to Figure 5.

Figure S6

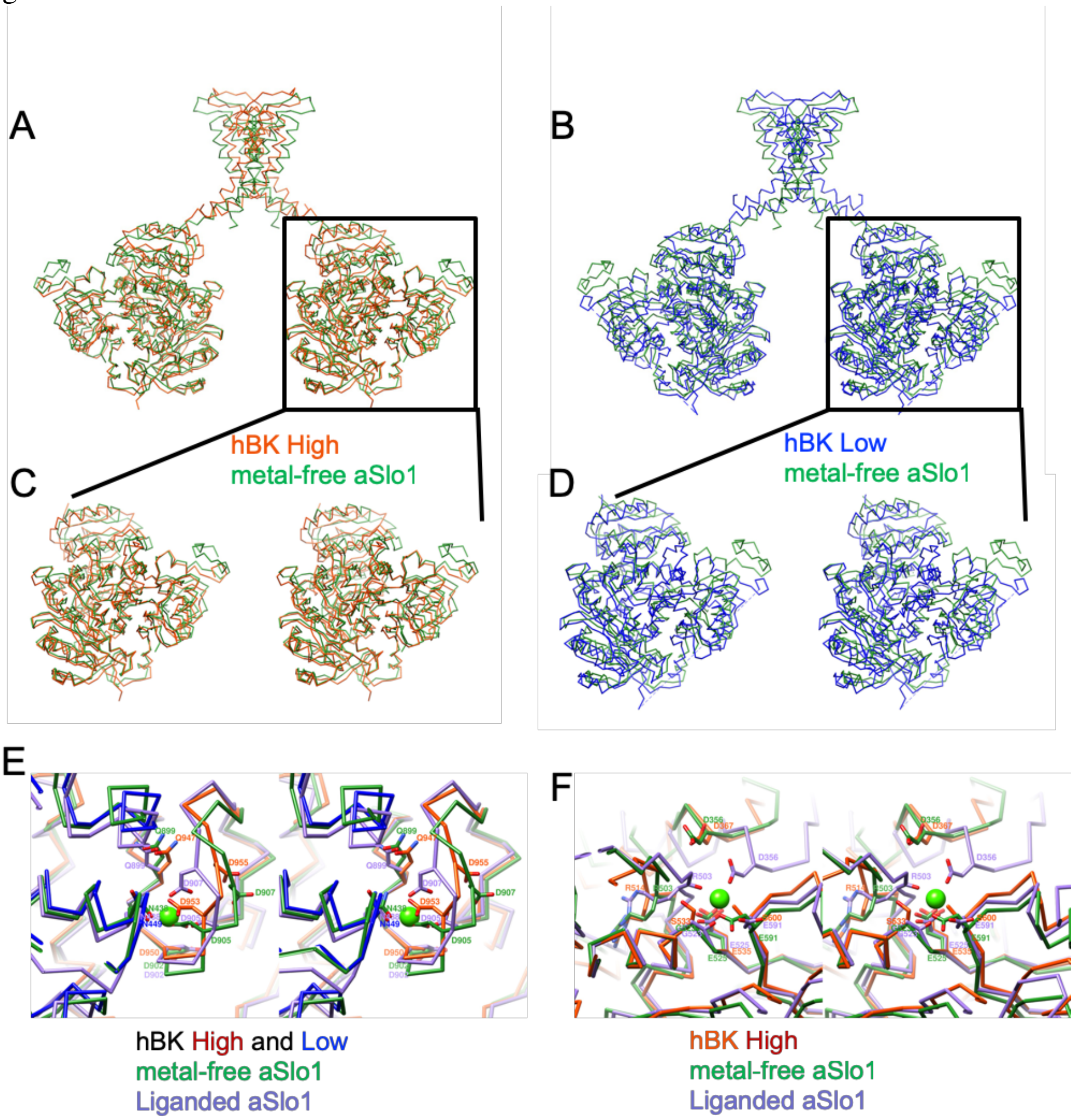


Figure S6: Comparison between hBK and aSlo1 (PDB: 5tji) structures. (A) Superposition of hBK High subunits (orange red) and two opposing aSlo1 subunits (green). (B) Superposition of hBK Low subunits (blue) and two opposing aSlo1 subunits (green). (C-D) Stereo views of one subunit shown in A and B respectively. (E) Comparison of the $Ca^{2+}$ bowl in metal-free hBK (orange red and blue), metal-free aSlo1 (green) and liganded aSlo1 (purple). (F) Comparison of the RCK1 $Ca^{2+}$-binding site in metal-free hBK, metal-free aSlo1 and liganded aSlo1. $Ca^{2+}$

coordinating residues are labeled, and the position of $Ca^{2+}$ ion in liganded aSlo1 was represented by a green sphere.

Related to Figure 2&4.
